# Impact of sucrose sinks on phloem transport

**DOI:** 10.1101/2025.01.28.635320

**Authors:** Mazen Nakad, Ensheng Weng, Pierre Gentine

**Affiliations:** Chemical Engineering Departmen, School of Engineering, Lebanese American University, Byblos, Lebanon; Center for Climate Systems Research, Columbia University, New York, NY, USA; NASA Goddard Institute for Space Studies, New York, NY, USA; Department of Earth and Environmental Engineering, Columbia University, New York, NY, USA; Center for Learning the Earth with Artificial Intelligence and Physics, Columbia University, New York, NY, USA

**Keywords:** M*ü*nch, Sucrose allocation, Mass flux, Taylor Dispersion

## Abstract

The movement of photosynthates within plants is a focus in plant physiology, ecohydrology, and earth systems modeling. The phloem, one of the plant’s hydraulic systems, facilitates this transport. It is believed to be optimized for efficient photosynthates transport, notably sucrose. This has implications ranging from local impacts on plant survival during drought to ecosystem-scale effects on carbon and water cycling. Most models for phloem transport rely on the pressure-flow hypothesis, where sucrose is loaded in leaves, drawing water from the xylem through osmosis, generating pressure gradients for transport. Experimental challenges in measuring sugar fluxes have led to reliance on theoretical models, though discrepancies exist, especially for long-distance transport. Criticism of the pressure-flow hypothesis notes low hydraulic conductance in sieve tubes, possibly hindering sucrose transport in taller plants. This research explores osmotically driven flows through the development of a new one-dimensional numerical model that includes sources from photosynthesis and sinks towards the stem and roots. The model also incorporates a concentration-dependent viscosity and the xylem water potential. It shows that different allocation schemes of sucrose sinks towards the stem of the plant influence the speed at which sucrose is transported. These findings provide insight into how carbon allocation along the phloem may have evolved to enhance the efficiency of transporting soluble compounds in the phloem.

**Highlights:**

- Sucrose sinks along the phloem influence mass flux beyond simple reaction-like consumption dynamics
- Mass flux is modulated by the sucrose allocation profile along the tree stem
- Observations of xylem-phloem water exchange can provide insights into the physical sucrose sink profile

## 1. Introduction

The transport of sucrose and other soluble organic compounds from photosynthesizing leaves to other parts of the plant for consumption or storage is a fundamental process in plant physiology. This process occurs via the phloem, a complex vascular tissue, and is primarily explained by the pressure-flow hypothesis (PFH), commonly referred to as the Münch mechanism. First proposed by Ernst Münch in the 1930s [1], the model remains a cornerstone of plant nutrient transport [2, 3]. According to the PFH, sugars are actively loaded into the phloem at the source (leaves), creating a high osmotic pressure that draws water into the phloem from the xylem. This water influx generates positive pressure, driving the bulk flow of the sugar solution through the sieve tubes to sinks (e.g., roots or storage organs) [4, 5, 6]. The Münch mechanism is widely accepted due to its simplicity and robustness in modeling phloem hydrodynamics [7, 8, 9, 10, 11, 12, 13, 14].

However, the PFH faces challenges, particularly for long-distance transport in large trees. One key critique is that the hydraulic conductance of sieve tubes may be insufficient to support long-distance transport solely via osmotic pressure-driven flow [15, 16, 17, 18, 19]. Experimental data suggest that the pressure gradients required would necessitate sucrose concentrations higher than observed, especially in tall trees [20, 21, 22]. This raises questions about the adequacy of the Münch mechanism for explaining phloem transport in large plants [15, 2, 23, 24, 25], especially considering that sieve plates, which are specialized structures within the phloem that separate sieve tube elements and contain pores through which sap flows, introduce additional resistance, further complicating the assumption of straightforward pressure-driven flow [10, 26, 27, 28, 29].

Alternative mechanisms have been proposed. One hypothesis suggests that transport involves a series of relay stations where sugars and water are exchanged along the phloem pathway, reducing the need for large pressure gradients [24]. This may improve transport efficiency in tall trees by providing local pressure regulation. Additionally, water exchange between the phloem and surrounding tissues, especially under stress, likely plays a significant role [30, 31, 32, 6]. This interaction allows phloem transport to adapt to varying environmental conditions and sap viscosities, enhancing its overall efficiency and resilience [33, 31, 6]. Emerging research on solute dynamics highlights the role of fluid viscosity in determining transport efficiency. Variations in sucrose concentration affect sap viscosity, which alters flow rates. Models accounting for viscosity-dependent flow have improved predictions, especially under conditions like drought, when sap composition changes [11, 10, 34, 35, 36]. Taylor dispersion, where solutes spread via diffusion and flow, may accelerate sucrose transport even without high-pressure gradients [37, 38], underscoring the importance of both radial and axial flows for efficient long-distance transport [36]. Structural aspects of the phloem, particularly sieve plates, are also under renewed investigation [29, 39, 40, 41, 26, 42]. Once thought to obstruct flow, sieve plates may maintain structural integrity, preventing collapse and supporting flow stability [43, 44, 45, 28]. In tall trees, sieve plates may buffer pressure fluctuations and help sustain long-distance transport [46].

Phloem transport efficiency has broader implications beyond plant physiology. Sucrose transport influences carbon allocation, plant health, and productivity [47, 48]. Phloem transport is also crucial to ecosystem-level processes, such as carbon and water cycling, vital for modeling plant responses to climate change [49, 50, 51]. Furthermore, hydraulic interactions between phloem and xylem are key to plant responses to environmental conditions [52, 53, 54, 55]. While Münch’s hypothesis focused on osmotic gradients, recent research emphasizes the importance of xylem-phloem water exchange, especially under stress [56]. To simulate these dynamics, several models have been developed to capture water movement and hydraulic coupling between xylem and phloem [30, 57, 58, 59, 60]. These models highlight the significance of hydraulic connectivity in supporting phloem transport, even under fluctuating environmental conditions. Tight coordination between the leaf-xylem-phloem network is fundamental to regulating growth, survival, and stress responses [49, 50, 47].

In this context, the present study builds on previous work by incorporating not only hydraulic connectivity but also the influence of sucrose sinks along the phloem pathway. Trees, particularly in large ecosystems, rely on these sinks for essential functions such as growth, embolism recovery, and defense against biotic threats like bark beetles. However, these sinks place additional demands on the phloem transport system, significantly altering the dynamics of sugar transport. This study develops a mathematical model that integrates local sucrose sinks along the phloem pathway. Additionally, xylem background pressure is incorporated into the PFH framework as one-way coordination, where the xylem water potential, sink demand, and photosynthesis are externally imposed using calculations from a demographic vegetation model. The primary focus of the present work is to examine the impact of the sucrose sink function, particularly its profile along the phloem pathway, on the speed of mass flux.

### 2. Theory

The basic equations governing sucrose transport in plants are first reviewed. The system is modeled as a water reservoir of length *L* (i.e., the xylem) surrounded by a thin layer of thickness *a* (i.e., the phloem). A longitudinal section of this system is illustrated in Figure 1. The phloem is described using a Cartesian coordinate system, where *z* = 0 is at the leaf level and *z* = *L* is at the root level in the longitudinal direction, while *y* = 0 marks the boundary between the xylem and the phloem, extending to *y* = *a* at the tree bark. In this framework, sucrose transport is modeled for a two-dimensional thin layer of the phloem, assuming a uniform distribution of sucrose in the angular direction. Although this is not a perfectly accurate mathematical representation, it simplifies the derivation by using Cartesian rather than radial coordinates. Consequently, the dynamic scaling of flow variables is influenced by the slender geometry, characterized by an aspect ratio *ϵ* = *a/L*⪡ 1.

**Figure 1:**
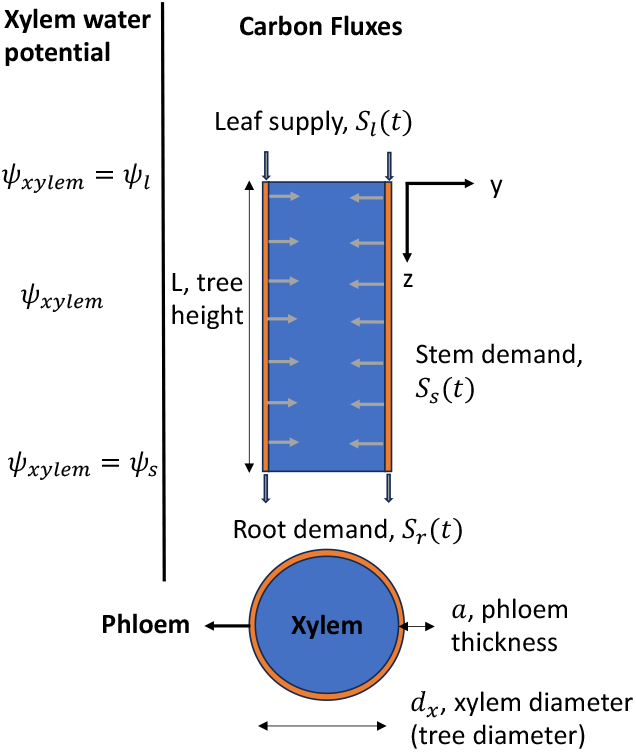
Schematic diagram of the xylem-phloem hydraulic system with carbon (sucrose) fluxes into and out of the phloem.

Sucrose fluxes into and out of the phloem, as well as the xylem water potential, are calculated using the demographic vegetation model BiomeE [61] and serve as external inputs for the model that solves transient phloem hydrodynamics between two time-steps. BiomeE provides hourly vegetation carbon and water fluxes, creating a one-way coupling from the root-xylem-leaf system to the phloem. The boundary between the xylem and phloem is modeled as a semi-permeable membrane with uniform permeability *k*, allowing the exchange of water molecules, but not sucrose molecules, between the two tissues.

In this model, the variables representing sucrose mass, fluid dynamic pressure, and the velocity components in the longitudinal and radial directions are denoted by *c, p, u*, and *v*, respectively, while the xylem water potential is represented by *ψ*. The relatively small longitudinal velocity *u* in plants allows for the approximation of flow as a low Reynolds number flow, where the Reynolds number is given by *Re* = *ρau/*⪡ 1. Here, *µ* is the dynamic viscosity and *ρ* is the water density, assumed independent of *c*, while dynamic viscosity can influence flow dynamics [36]. This approximation permits the neglect of inertial effects in the longitudinal momentum equation, with viscous forces becoming dominant. Additionally, frictional losses from sieve plates in the phloem are disregarded in the model, although such losses could be significant in some cases [21]. Despite these simplifications, the model offers a practical basis for exploring the effects of the spatial distribution of sucrose consumption in the stem on mass flux. For mass transport, the relative contributions of advective versus diffusive transport are quantified by the Peclet number, *Pe* = *au/D*, where *D* is the molecular diffusion coefficient, assumed independent of *c*. The Peclet and Reynolds numbers are related through the Schmidt number, *Sc* = *Pe/Re*, which for sucrose in water is large (*>* 10, 000). This suggests that advective transport plays a significant role in the solute mass balance, in contrast to the momentum balance, where inertial effects are negligible.

### 2.1 Conservation equations

The derivation of the governing equations for the flow field in two dimensions with concentration-dependent viscosity can be found in the literature [36]. Although a Cartesian coordinate system is used in this model, the conservation laws remain the same, leading to the following non-dimensional forms of the governing equations:

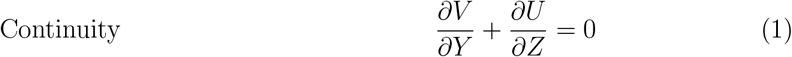

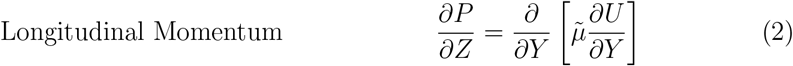

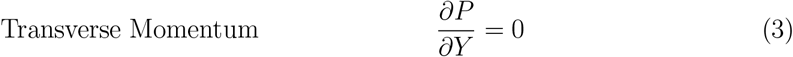

In these equations, the normalized variables are embedded in the formulation, where *u* = *u*_0_*U*, *v* = *v*_0_*V*, *p* = *p*_0_*P*, *y* = *aY* and *z* = *LZ*. with *u*_0_, *v*_0_, and *p*_0_ are characteristic axial velocity, radial velocity, and pressure. The characteristic length scales in the longitudinal and transverse dimensions are the phloem length *L* and thickness *a*. The longitudinal velocity scale *u*_0_ = *v*_0_*/ϵ* is determined from the continuity equation, and *p*_0_ = (*Lµ*_0_*u*_0_)*/a*^2^ is the viscous pressure scale.

These equations represent the leading-order approximation of the flow field. When the reduced Reynolds number tends to zero (i.e., *ϵ Re* → 0, as in lubrication theory) and the Schmidt number is sufficiently large (*Sc*⪢1), inertial forces and certain components of the tangential stress can be neglected. In this case, the flow field can be approximated as steady-state, where the advective forces of sucrose molecules dominate over the inertial forces of water molecules. As a result, the pressure field becomes dependent only on the longitudinal direction. A more general treatment can be found elsewhere [62]. Further simplifications can be made by assuming that viscosity variations in the transverse direction are negligible. This assumption leads to the following flow field equations:

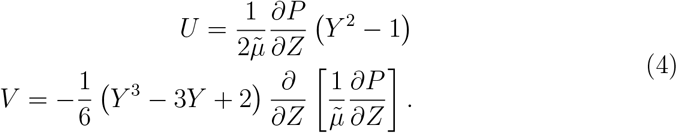

This simplification is critical for mathematical tractability and is supported by the fact that sucrose mass variations in the transverse direction are negligible compared to the area-averaged sucrose mass profile, as discussed later. Equations 4 were derived by applying the no-slip boundary conditions at *Y* = 0 and *Y* = 1 for the longitudinal velocity *U*, and the no-flow boundary condition at *Y* = 1 for the transverse velocity *V* . For completeness, these equations are also expressed in dimensional form as

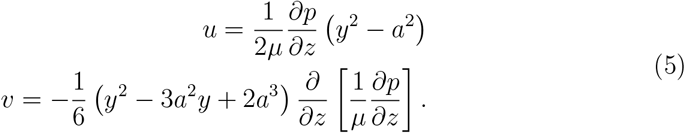

Equations 4 (or 5) describe the flow field for osmotically driven flows, neglecting variations in dynamic viscosity in the transverse direction. If one assumes that the dynamic viscosity is independent of sucrose concentration,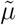 effectively becomes equal to one, simplifying the problem to a Hagen-Poiseuille formulation with a concentration-dependent pressure gradient, *∂p/∂z*, driven by osmosis.

However, since these flow field equations are incomplete, with two equations for three unknowns (*p, v*, and *u*), an additional boundary condition for *v* at *y* = *a* is required. This boundary condition must be provided by Darcy’s law and osmoregulation [63, 64]. The water flow between the xylem and phloem is driven by the water potential difference between these two hydraulic systems, and the non-dimensional form of the boundary condition is written as:

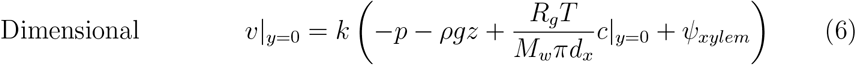

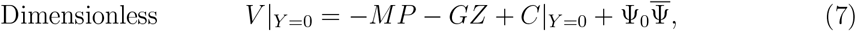

where *M* = *kµ*_0_*L*^2^*a*^*−*3^ is the Münch number, *G* = *ρgLM*_*w*_*πd*_*x*_(*R*_*g*_*Tc*_0_)^*−*1^ is a nondimensional number reflecting the relative importance of gravitational forces over osmotic potential, Ψ_0_ = *ψ*_*l*_*M*_*w*_*πd*_*x*_(*R*_*g*_*Tc*_0_)^*−*1^ reflects the xylem water potential’s importance relative to osmotic potential. Here *g, M*_*w*_, *d*_*x*_, *R*_*g*_, *T*, *c*_0_, and *ψ*_*l*_ represent the gravitational constant, sucrose molecular weight, xylem diameter, ideal gas constant, temperature, characteristic sucrose concentration, and leaf water potential (or xylem water potential scale), respectively. The transverse velocity scale, *v*_0_ = *kR*_*g*_*Tc*_0_(*M*_*w*_*πd*_*x*_)^*−*1^, is embedded in the formulation with *k* with representing membrane permeability. By applying this boundary equation to the transverse velocity equation, the flow field can now be solved simultaneously with the evolution of sucrose mass (discussed later). This results in four equations with four unknowns (excluding the dynamic viscosity, which is related to sucrose concentration/mass and adds an additional equation).

The conservation of sucrose mass is derived using the Reynolds transport theorem. The transport of sucrose in both longitudinal and transverse directions is governed by advection and molecular diffusion. The solute mass balance equation can be written as:

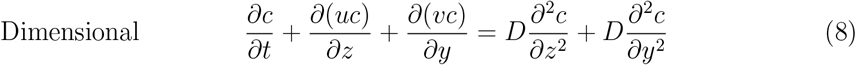

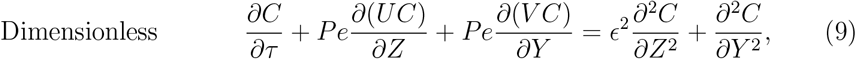

where the characteristic timescale *t*_0_ = *a*^2^*/D* is used for the time variable *t* = *τt*_0_, determined by the transverse molecular diffusion term, as it is the fastest process in the system. Along the longitudinal direction, sucrose enters the phloem from the leaf and exits towards the root:

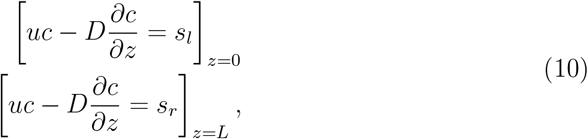

where *s*_*l*_ and *s*_*r*_ are the source and sink terms at the leaf and root, respectively. Along the transverse direction, a sucrose sink towards the stem at *y* = 0 and a no mass-flux condition at *y* = *a* are imposed:

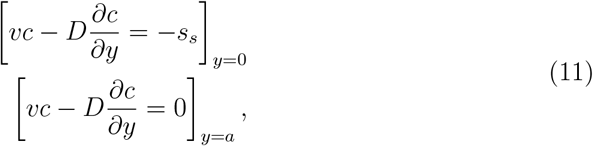

where *s*_*s*_ represents the stem sink, which can vary along the longitudinal direction since several factors, including structural demand, safety, or growth, influence sucrose demand in the stem. Therefore, multiple sink profiles will be employed to study the impact of this formulation on mass flux. At this point, the characteristic sucrose concentration scale *c*_0_ remains undefined. To define this scale, one can either base it on the sources and sinks (by scaling the boundary conditions and relating *c*_0_ to any of the sucrose flux terms *s*_*·*_) or derive it from the xylem-phloem water balance, scaling it with the xylem water potential. Physically, it makes more sense to scale *c*_0_ according to the more limiting factor. However, due to the generality of the problem, *c*_0_ will be scaled with the xylem water potential leading to *c*_0_ = *ψ*_*l*_*M*_*w*_*πd*_*x*_(*R*_*g*_*T*)^*−*1^ (i.e. Ψ_0_ = 1). This implies that the osmotic potential is balanced by the xylem water potential, and the transverse inflow of water is driven by the background pressure from the xylem. This scaling approach is similar to that adopted by [65], which balances osmotic pressure with xylem water potential and phloem pressure (via the relation *v*_0_ = *ϵu*_0_). In the limit of water potential equilibrium (large *M*), all variables are rescaled by *M*, leading to an approximate equality between the phloem water potential (including dynamic, hydrostatic, and osmotic components) and the xylem water potential. This occurs because the term on the left-hand side, *V/M*, approaches zero as *M* becomes large.

### 2.2. One-dimensional model for sucrose transport in plants

To arrive at the one-dimensional model that includes a stem sucrose sink along the phloem pathway, one can apply the area-averaging method of [37], with the only difference being the use of a Cartesian coordinate area-averaging operator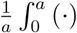 d*y* instead of the cylindrical one used in that method. In this case, the stem sink term appears after averaging equation 8, as a result of applying the boundary condition of equations 11. Only the key steps and differences will be presented here; for a full derivation of the method, please refer to [37].

Starting from equation 8, the one-dimensional model for sucrose transport is derived by using the area-averaging operator and imposing the boundary conditions from 11, leading to the following result:

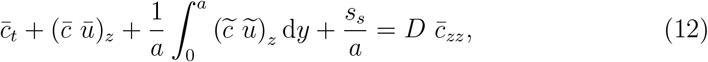

where 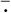 and 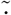denote area-averaged and deviation components respectively. Here, the integral term represents the Taylor dispersion contribution, and the last term on the left-hand side defines the sink term, which can be dependent on *z, t*, and *c*. As a starting point, only the position dependence will be studied here. Time dependence will not affect the model derivation and is not expected to play a significant role in phloem applications, where sucrose growth and storage change predominantly over longer timescales (e.g., daily, seasonal, or yearly). However, time dependence may become important during extreme events, such as bark beetle attacks, where rapid sucrose mobilization or shifts in demand could occur on shorter timescales due to heightened defensive or metabolic needs. In contrast, concentration dependence will influence the model derivation (except in the linear case where *s*_*s*_ *~ c*), since a concentration-dependent function *s*_*s*_ *~ f* (*c*) cannot be approximated by the area-averaged concentration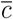 (i.e.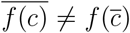. Unless the function is already specified in terms of area-averaged quantities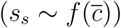, this adds complexity to the problem and is better left for future investigation.

The only unknown in equation 12 is 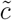, which requires an ordinary differential equation (ODE) that can be obtained by deriving the ‘deviation’ equation and applying dominant balance argument, leading to:

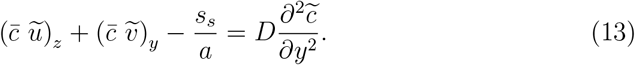

Here, the dominant balance is similar to that discussed in [37], with the addition of the sink term, which scales as 𝒪(1*/a*). Now,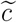 can be approximated by solving this ODE, applying the boundary condition at *y* = *a* where 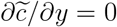(since *v* = 0 at *y* = *a* and 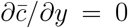 by definition), and recognizing that the area-averaged concentration deviation is zero by definition. This leads to the following result

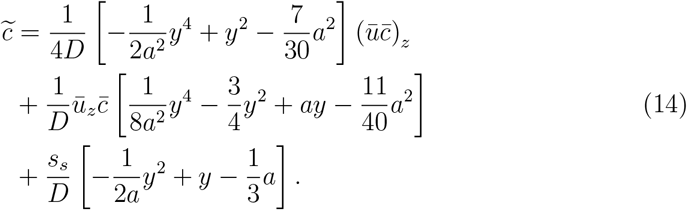

Using the result of equation 14 in combination with the flow field information from equations 5, one can derive the area-averaged one-dimensional model for sucrose transport with a continuous sink:

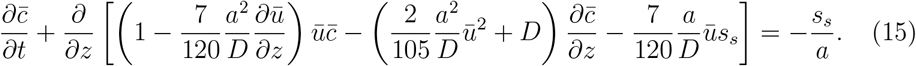

Several key differences between this result and the one developed in [37] should be noted. First, the sign of the advective term due to Taylor dispersion is negative. This arises from the way the problem was formulated. Although the two formulations appear contradictory, the difference is related to 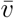, which is driven by osmosis. In this case, 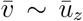, while in [37] 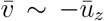. When expressed in terms of 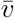 (the true ‘physical’ term), the sign is consistent with the previous work. Second, the coefficients differ due to the change in a coordinate system. Finally, a new term emerges due to the impact of the sink term. Physically, Taylor dispersion approximates transverse variations in concentration into area-averaged quantities for the one-dimensional equation. Since the stem sink term directly affects the transverse direction (acting as a boundary condition that alters the concentration profile), Taylor dispersion naturally introduces an additional term. This term appears as a flux term that slows down sucrose propagation. A reduced amount of sucrose decreases osmotic potential, leading to a lower transport speed.

The final step in this derivation is to ensure mass balance. To achieve this, the boundary conditions in equations 10 need to be rewritten in terms of the area-averaged components, leading to:

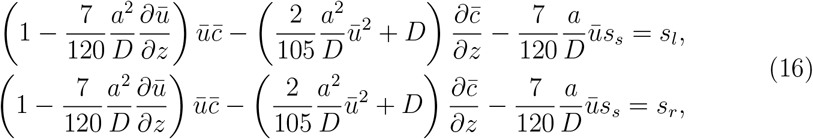

where the first equation defines the boundary condition at the entrance (leaf level), and the second equation defines the boundary condition at the exit (root level). If one integrates equation 15 along the longitudinal direction and applies the boundary conditions from equations 16, the evolution of total mass in the phloem is governed by the following ODE:

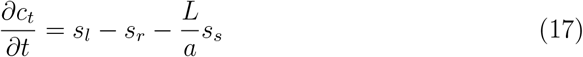

where *c*_*t*_ is the total sugar in *grams/meter* (since the problem is defined on a thin layer of the phloem in the angular direction, multiplying by *πd*_*x*_ gives the total sucrose in the phloem). The terms *a* and *L* appear due to the area-averaging and integration process: *a* results from averaging, representing the characteristic transverse length scale of the domain over which the boundary sink term influences the local concentration, and *L* results from integration, effectively transforming a pointwise sink into a global sink that represents the total sucrose removal along the entire length of the phloem. A description of the variables and their respective unit is shown in table 1. The numerical method used to simulate phloem hydrodynamics with the necessary initial and boundary conditions and the inclusion of a concentration-dependent viscosity are described in Appendix A.

**Table 1:**
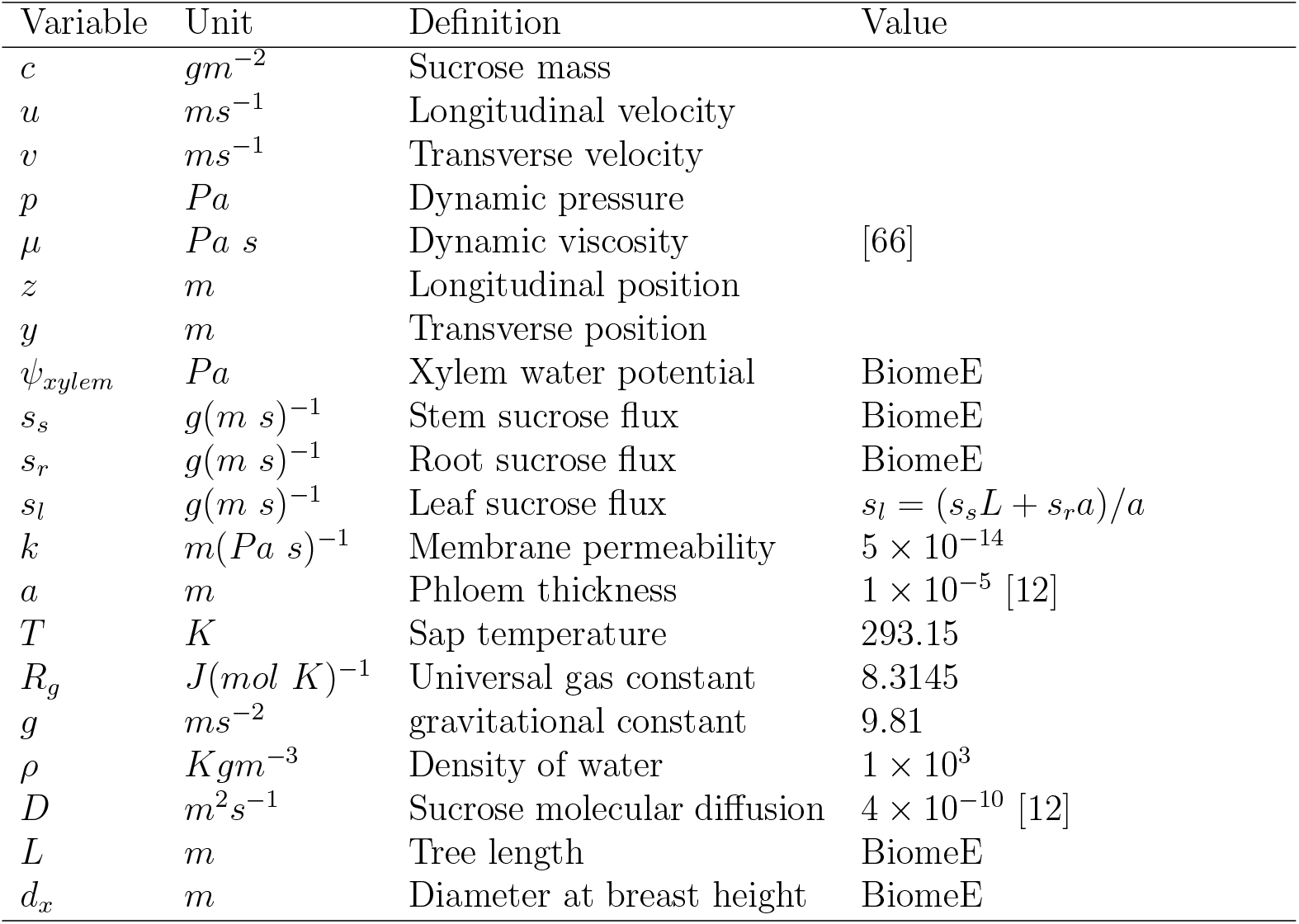
Summary of variables, units, and corresponding values or sources.

| Variable | Unit | Definition | Value |
| --- | --- | --- | --- |
| $c$ | $gm^{-2}$ | Sucrose mass | |
| $u$ | $ms^{-1}$ | Longitudinal velocity | |
| $v$ | $ms^{-1}$ | Transverse velocity | |
| $p$ | $Pa$ | Dynamic pressure | |
| $\mu$ | $Pa\ s$ | Dynamic viscosity | [66] |
| $z$ | $m$ | Longitudinal position | |
| $y$ | $m$ | Transverse position | |
| $\psi_{xylem}$ | $Pa$ | Xylem water potential | BiomeE |
| $s_s$ | $g(m\ s)^{-1}$ | Stem sucrose flux | BiomeE |
| $s_r$ | $g(m\ s)^{-1}$ | Root sucrose flux | BiomeE |
| $s_l$ | $g(m\ s)^{-1}$ | Leaf sucrose flux | $s_l = (s_s L + s_r a)/a$ |
| $k$ | $m(Pa\ s)^{-1}$ | Membrane permeability | $5 \times 10^{-14}$ |
| $a$ | $m$ | Phloem thickness | $1 \times 10^{-5}$ [12] |
| $T$ | $K$ | Sap temperature | 293.15 |
| $R_g$ | $J(mol\ K)^{-1}$ | Universal gas constant | 8.3145 |
| $g$ | $ms^{-2}$ | gravitational constant | 9.81 |
| $\rho$ | $Kgm^{-3}$ | Density of water | $1 \times 10^3$ |
| $D$ | $m^2s^{-1}$ | Sucrose molecular diffusion | $4 \times 10^{-10}$ [12] |
| $L$ | $m$ | Tree length | BiomeE |
| $d_x$ | $m$ | Diameter at breast height | BiomeE |

### 2.3 Position dependent stem demand

As discussed in section 2.2, the stem demand can be time, position, or concentration dependent. Time dependence arises due to external factors, such as extreme events where a higher demand for sucrose is needed for a short period, often for safety. For longer processes, such as growth, the sink demand can be assumed to be time-independent, as the plant stores sucrose in a steady-state manner to be utilized over a longer timescale. This storage and use are part of the plant’s overall carbon balance, which is influenced by various physiological and ecological factors but lies outside the scope of this study.

Concentration dependence is related to the underlying physical processes by which the plant stores sugar, such as reaction-like processes, diffusion, or advection. In this case, the dependence of the sink on concentration can be linear or nonlinear, which could affect the derivation of the model. Other assumptions might be required to maintain mathematical simplicity. Since the primary processes by which plants store sugar are still under investigation, and the main focus of this study is on the impact of longitudinal variations in sucrose demand along the stem, concentration dependence will not be tackled here.

The main focus of this study will be on position dependence, which can arise from the structural behavior of the plant. For instance, the bottom of the stem may have a thicker bark, while new branches at the top are still sucrose-starved (i.e., photosynthesis at this level does not produce enough sucrose to meet the structural demands, so sucrose is supplied from the source regions at the top of the tree, where sucrose levels are higher). In this case, different functions for the sink can be specified based on the location in the phloem.

In this study, five functions will be explored:

- Constant sink (‘CS’): where the sink is independent of position, given by *s*_*s*_ = (*f*_*g*_ +*f*_*r*_)*L*^*−*1^ where *f*_*g*_ is the stem growth demand, and *f*_*r*_ is the stem respiration (growth and maintenance respiration)
- Linear sink highest at bottom (‘LSb’): *s*_*s*_ = 2(*f*_*g*_ + *f*_*r*_)*zL*^*−*2^
- Linear sink highest at top (‘LSt’): *s*_*s*_ = 2(*f*_*g*_ + *f*_*r*_)(*L − z*)*L*^*−*2^
- Linear growth sink highest at top (‘LGSt’): *s*_*s*_ = 2*f*_*g*_*zL*^*−*2^ + *f*_*r*_*L*^*−*1^
- Linear growth sink highest at bottom (‘LGSb’):*s*_*s*_ = 2*f*_*g*_(*L − z*)*L*^*−*2^ + *f*_*r*_*L*^*−*1^

In the cases of ‘LGSb’ and ‘LGSt’, the variation between respiration and growth demand reflects the underlying biological processes: respiration is linked to the plant’s current metabolic state, while growth demand accounts for the allocation of resources for future development. These formulations capture this distinction by separating the immediate energy needs (respiration) from the long-term resource allocation for growth. Higher-order polynomials can also be formulated but would complicate the analysis, which focuses on how spatial variations affect the flow speed. When integrated over the longitudinal direction, all these functions yield *s*_*s*_ = *f*_*g*_ + *f*_*r*_ which is predefined by BiomeE.

## 3. Results

### 3.1. Impact of steady-state assumptions on phloem hydrodynamics

Many studies on phloem sucrose transport assume a steady-state condition, where sucrose fluxes are constant, and there is no significant sucrose storage within the phloem. This assumption is reasonable because the phloem’s volume is negligible compared to that of the stem and roots. In such cases, the fluxes into the phloem (from photosynthesis) and out of the phloem (to the stem and roots) are balanced and much larger than the small amount of sucrose stored within the phloem flow. However, in real plants, photosynthetic rates vary significantly over the 24-hour diurnal cycle. If the stored starch in leaves does not compensate for these variations, the differences between the source (leaves) and sink (roots and stem) may induce transient behavior in the phloem. This is especially relevant for long-distance transport, where the time required for sugars to travel from source to sink can range from a few hours to as long as 9 hours, depending on the tree height and environmental conditions. Moreover, under extreme conditions, such as drought, changes in xylem water potential may alter the amount of sugar required to maintain effective phloem transport. These changes could also affect sugar demand in the stem, where sugars might be used for processes like cell repair after cavitation events.

Figure 2 compares steady-state phloem dynamics (where the time derivative in equation 15 is zero), labeled ‘SS’, with two transient models: one using an initial guess for *α* = 2, described in Appendix A.3, labeled ‘TRI’, and another with steady-state initial conditions, where transient behavior evolves over time, labeled ‘TRS’. These models assume a uniform stem sink (‘CS’ from section 2.3). The results depict the flow behavior over a 24-hour cycle for a tree in equilibrium under normal environmental conditions (using BiomeE) and at the final phloem time step. For example, if *dτ* = 0.1 and *n*_*t*_ = 144000, it corresponds to the 1-hour time step imposed by BiomeE, and the results are displayed at this value of *n*_*t*_.

**Figure 2:**
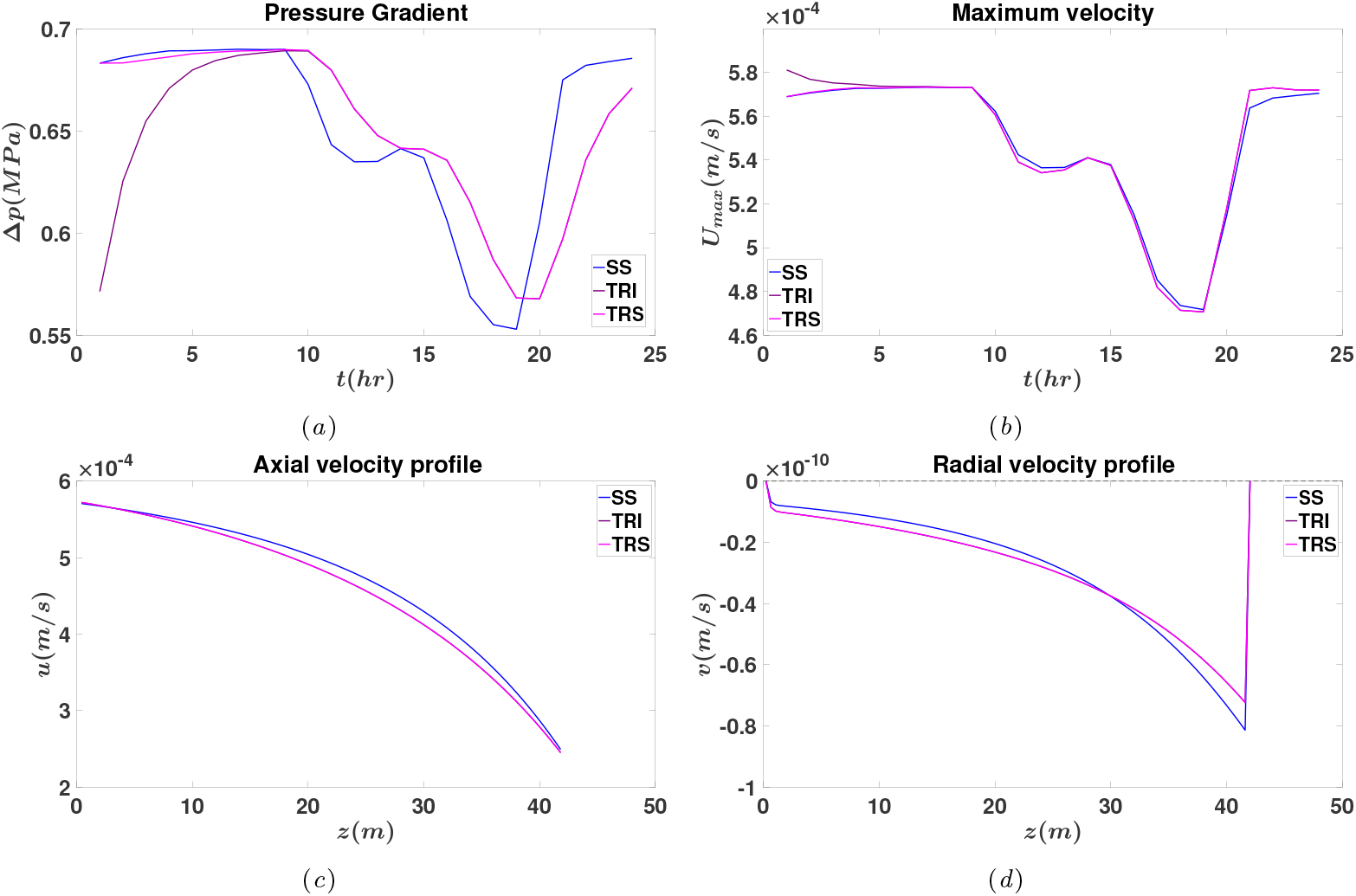
Results from a 24-hour cycle simulation showing the behavior of different models: steady-state (SS) in blue, transient phloem dynamics with an initial guess using *α* = 2 (TRI) in purple, and transient dynamics with steady-state initial conditions (TRS) in pink. (a) Pressure gradient evolution over time, (b) maximum phloem velocity as a function of time, (c) longitudinal velocity profile at the final time step, and (d) transverse velocity profile at the final time step.

The results suggest that the steady-state assumption is generally valid under normal, non-stressed environmental conditions. The only notable difference appears in Figure 2a, which shows the pressure gradient as a function of time. In the ‘TRI’ case, the pressure gradient (calculated as Δ*p* = *p*_1_*−p*_*n*_) takes a few hours to stabilize, making it more susceptible to the diurnal variations in xylem water potential (not shown but similar to the ‘SS’ case). This behavior is expected because the xylem water potential is calculated under a steady-state assumption over the hour and then imposed on the phloem dynamics as a forcing condition. Consequently, any rapid changes in xylem water potential impact the phloem at a slower pace. However, in real plants, xylem and phloem dynamics are interdependent, with changes in xylem water potential sensed by the phloem in real time. If the xylem hydrodynamics were solved at smaller intervals (e.g., around 1 second), these transient effects would not be as pronounced.

Figure 2b shows the maximum velocity in the phloem over time. Here, all cases exhibit similar behavior since the maximum velocity is influenced not only by the xylem water potential and phloem pressure gradients but also by the sink boundary conditions and the uniform stem sink, which evolve concurrently. The differences were even less significant for the total sucrose mass and the integrated phloem water potential (both not shown), as these quantities are closely tied to the imposed boundary conditions on the phloem. Figures 2c and 2d display the longitudinal and transverse velocity profiles in the *z* direction at the final time step, respectively. From these figures, it is evident that the differences between the cases are negligible. This trend was also observed for the sucrose mass and pressure profiles (not shown).

### 3.2. Effect of longitudinal variations in dynamic viscosity on phloem transport

The transport of sucrose within the phloem from the leaf to the roots occurs at low Reynolds numbers, indicating a balance between the pressure gradient and frictional losses along the phloem. Both of these forces are concentration-dependent. The pressure gradient arises from a sucrose gradient, which translates into an osmotic potential difference between the source and the sink, while frictional losses are influenced by shear stresses, which in turn depend on sucrose concentration through the dynamic viscosity *µ*. An increase in the sucrose gradient introduces an additional forcing term in the transverse momentum equation 4, beyond the local effects of viscosity (or a reduction in the phloem’s hydraulic conductivity) in the longitudinal momentum equation 4. To investigate this effect, we compare a constant viscosity model, where viscosity is calculated using the sucrose characteristic scale *c*_0_ (the mean sucrose mass required to maintain the osmotic potential needed to extract water from the xylem), with a ‘variable’ viscosity model that accounts for local variations and gradients in viscosity. Figure 3 presents results for three different stem sink profile cases (‘CS’, ‘LST’, and ‘LGST’).

**Figure 3:**
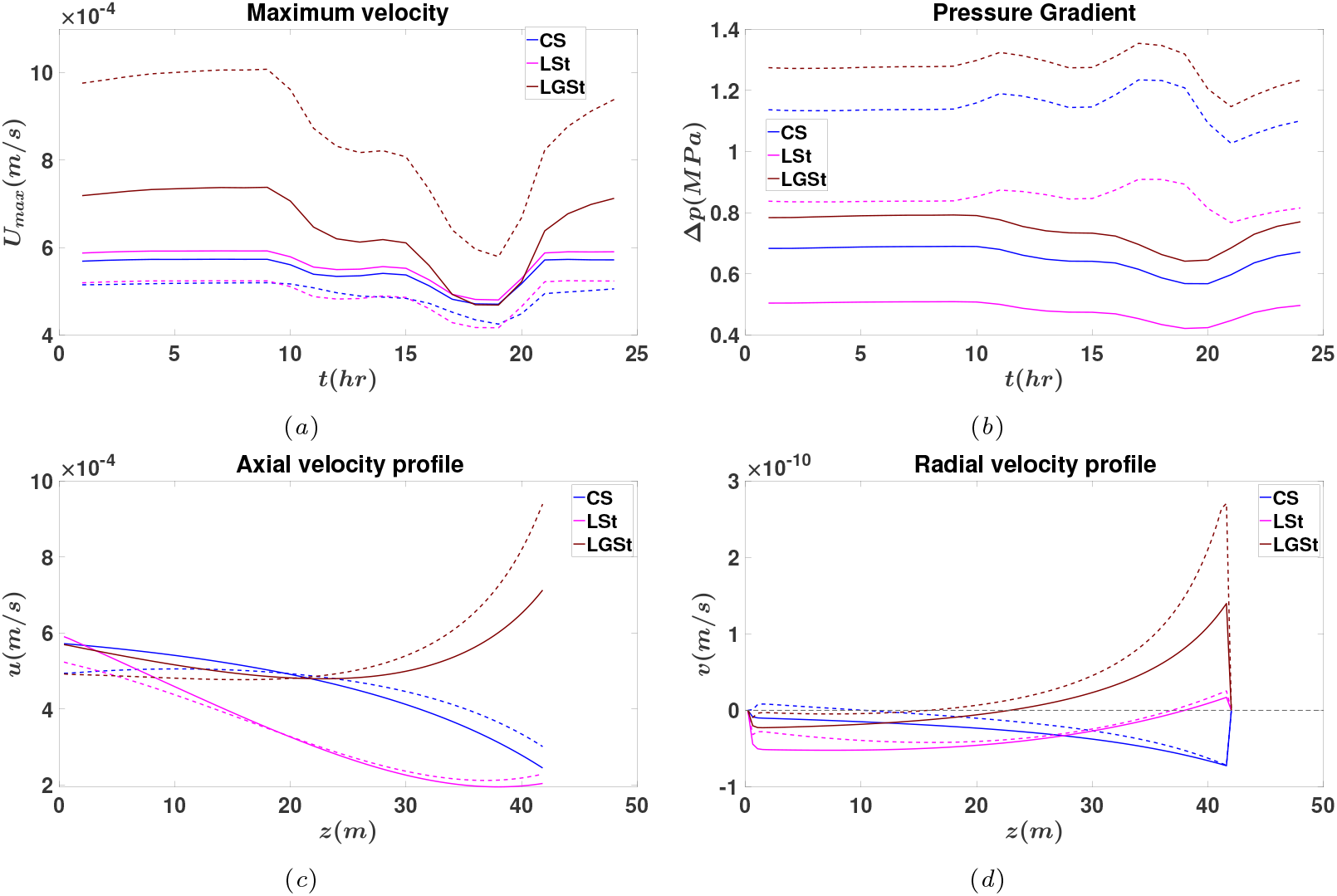
Results from a 24-hour cycle simulation showing the behavior of two models: variable viscosity (dashed lines) and constant viscosity (solid lines). Different colors represent different sink profiles: constant sink (‘CS’) in blue, linear growth and respiration with high sink at the top (‘LSt’) in pink, and linear growth with a high sink at the top (‘LGSt’) in brown. (a) Maximum velocity of the phloem flow, (b) pressure gradient across the phloem, (c) longitudinal velocity profile at the final time step, and (d) transverse velocity profile at the final time step.

Figure 3a shows the evolution of the maximum longitudinal velocity during the diurnal cycle. In this figure, only the case with linear growth and a high sink at the top (‘LGSt’) shows a significant increase in velocity compared to the constant sink (‘CS’) and the linear growth with respiration and a high sink at the top (‘LSt’). However, this increase can be misleading, as the sink variations along the phloem also affect the velocity profiles, as shown in Figures 3c and 3d, and discussed further in 3.3. From Figure 3c, it is evident that the maximum velocity along the phloem depends on the imposed sink profile. Despite this, the effect of viscosity on flow is consistent across all cases. At the beginning of the phloem, an increase in the sucrose required to satisfy the imposed inlet flux leads to a decrease in longitudinal velocity (due to reduced hydraulic conductivity). Conversely, near the outlet, a reduction in sucrose requirements results in an increase in longitudinal velocity (due to improved hydraulic conductivity).

Figure 3d highlights distinct behavior in the transverse velocity profile, especially for the ‘CS’ case. In the constant viscosity model, the transverse velocity is consistently negative, indicating water movement from the phloem into the xylem.

However, in the variable viscosity model, water flows into the phloem near the inlet and towards the xylem near the outlet. This behavior aligns more closely with the Münch mechanism, where water moves toward regions of high sucrose concentration (near the source) and flows towards the xylem in regions with lower sucrose concentration (near the sink). This suggests that incorporating dynamic viscosity provides a more accurate representation of natural water movement patterns in plants. In the other two cases (‘LGSt’ and ‘LSt’), a decrease in the magnitude of the transverse velocity near the inlet and an increase near the outlet is observed.

These variations are closely linked to the pressure gradient, as shown in Figure 3b. In all cases, an increased pressure gradient is evident due to elevated inlet pressure and reduced outlet pressure (not shown). The pressure gradient is influenced by the viscosity gradient, which adds a first derivative term to the osmosis equation 6 due to the relationship between velocity (*v*), viscosity (*µ*), and pressure (*p*) in the flow field equation 5. The inclusion of this term results in steeper pressure gradients near both the inlet and the outlet. In contrast, when this term is not included, the pressure profile becomes more diffusive, with smoother transitions across the phloem. Essentially, the viscosity gradient modifies how the pressure gradient is distributed along the phloem, influencing the velocity and flow direction. Despite these variations, the total sucrose mass remains similar across all cases (not shown), suggesting that these changes in the velocity profile do not significantly affect the overall sugar transport. The integrated phloem potential shows minimal change, with a slight decrease observed in all cases (not shown).

### 3.3. Impact of Stem Sink Variations Along the Phloem

This section examines the effect of stem sink variations on flow dynamics using the constant viscosity model with transient dynamics and steady-state initial conditions. While the variable viscosity model was explored in earlier sections, it is excluded here for two main reasons: (1) stem sink variations exhibit similar trends in both models for all cases except the constant sink case (‘CS’), as discussed and shown in Figure A.5; and (2) the constant viscosity model provides greater numerical accuracy and stability under the current simulation setup. For the variable viscosity model, increasing the grid resolution from *n* = 100 to *n* = 1000 resulted in a percentage error of approximately 5%. This error arises from the added nonlinearity introduced by the concentration-dependent viscosity, which affects the accuracy and stability of the analytically solved Jacobian matrix. In contrast, the constant viscosity model exhibited significantly lower error (less than 1%) under the same conditions, making it more robust for analyzing the influence of stem sink profiles. Addressing these challenges for the variable viscosity model, such as through advanced numerical schemes or using a numerical Jacobian with preconditioning, lies outside the scope of this study but represents an avenue for future improvement. By focusing on the constant viscosity model, this section highlights how different stem sink profiles influence flow speed and longitudinal concentration profiles in the phloem.

The simulation results for the constant viscosity model using the five sink profiles discussed earlier are shown in Figure 4. These results highlight how variations in stem sink profiles affect flow dynamics, particularly the flow speed and pressure gradients along the phloem. From Figure 4a, it is evident that the maximum velocity attained is influenced by the sink profile. Most cases exhibit similar trends over the 24-hour cycle, except for the ‘LGSt’ case, which shows a substantial decrease in velocity between hours 15 and 20. This period corresponds to the time when the xylem water potential is at its lowest, and sinks are at their peak (not shown). The decrease in velocity for the ‘LGSt’ case is directly related to its longitudinal velocity profile (Figure 3c), where the highest velocity is observed at the end of the phloem. Having the maximum velocity at the end of the phloem makes the system more vulnerable to external forcings, such as low xylem water potential and heightened respiration demands (near the top of the tree for this case). These additional stressors increase the resistance the phloem must overcome to maintain flow. In contrast, other sink profiles exhibit more balanced velocity distributions along the phloem, reducing their sensitivity to external perturbations.

**Figure 4:**
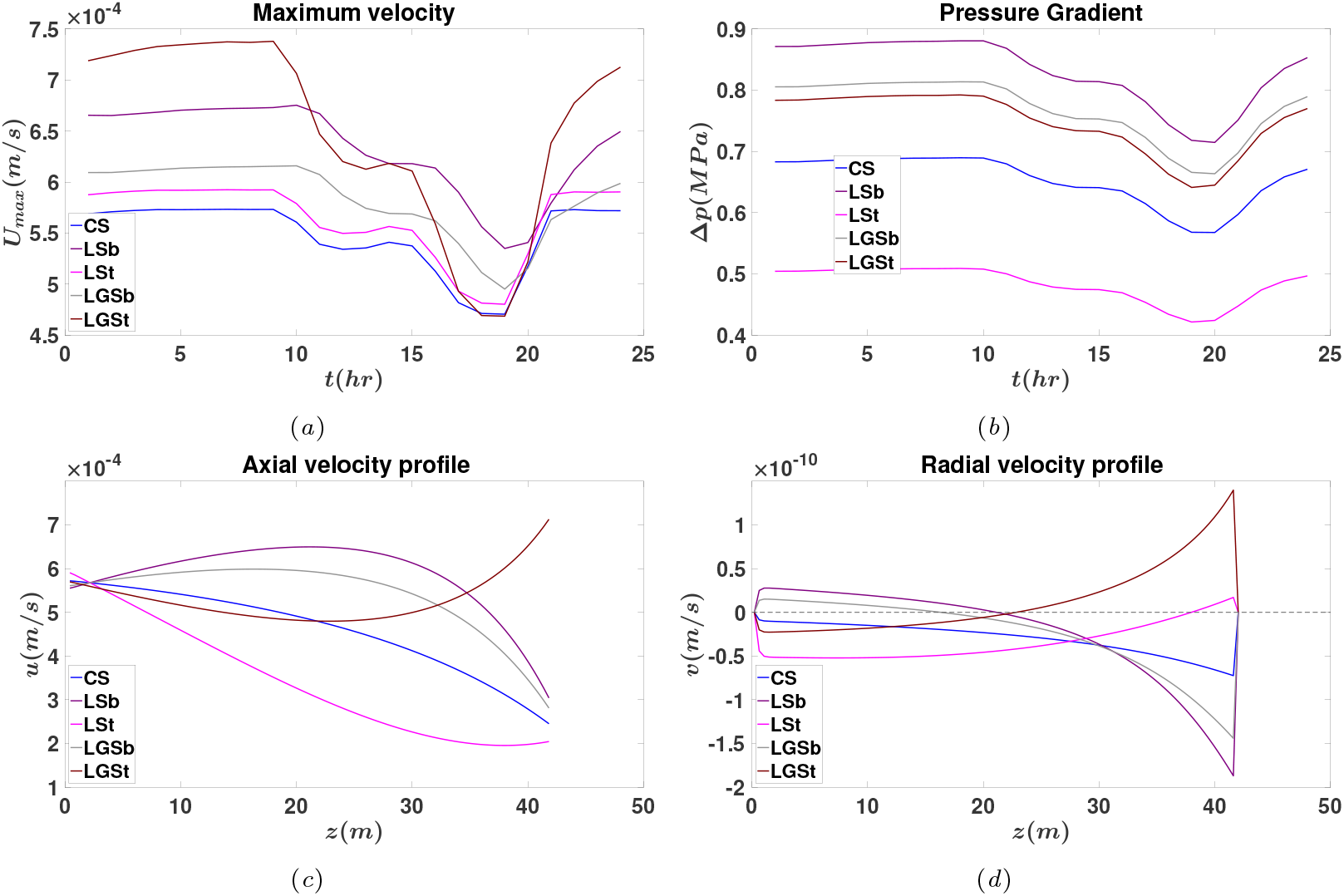
Results from a 24-hour cycle simulation showing the behavior of the constant viscosity model. Different colors represent different sink profiles: constant sink (‘CS’) in blue, linear growth and respiration with high sink at the top (‘LSt’) in pink, linear growth with a high sink at the top (‘LGSt’) in brown, linear growth and respiration with high sink at the bottom (‘LSb’) purpule and linear growth with a sink at the bottom (‘LGSb’) in grey (a) Maximum velocity of the phloem flow, (b) pressure gradient across the phloem, (c) longitudinal velocity profile at the final time step, and (d) transverse velocity profile at the final time step.

A clearer understanding of these dynamics is provided by the pressure gradient, shown in Figure 4b, which represents the driving force for flow, generated by osmosis to counteract the water potential gradient. Across all sink profiles, the pressure gradient exhibits similar trends over time, closely following external forcings such as the xylem water potential and the total stem and root sinks. This behavior reflects the integrated, large-scale dynamics of the phloem system. Among the sink profiles, the ‘LGSb’ case demonstrates the highest driving force, while the ‘LSt’ case exhibits the lowest. This disparity arises from how sink variations influence the osmotic potential along the phloem. In the ‘LGSb’ case, where the sink demand is smaller (accounting only for growth) and concentrated near the base, the osmotic potential at the source remains high, enhancing the driving force. In contrast, the ‘LSt’ case, with a larger sink demand (accounting for both respiration and growth) concentrated near the top, results in more rapid sucrose removal near the source, reducing the osmotic potential and thus diminishing the driving force. Interestingly, the ‘LGSt’ case shows an improvement in driving force compared to the ‘CS’ case, suggesting that allowing small variations in the sink profile at the top can enhance flow dynamics when sucrose transport is primarily driven by advection (*U* ≠ 0). Another important trend is the effect of sink placement. When sucrose is predominantly removed near the base of the phloem, as in the ‘LGSb’ case, the osmotic potential remains high along most of the pathway, sustaining the driving force. The distinction between the ‘LSb’ and ‘LGSb’ cases lies in the early phloem behavior: in the ‘LSb’ case, more sucrose is removed near the source, causing a slight reduction in osmotic potential compared to the ‘LGSb’ case, where sucrose removal is more gradual.

The transverse velocity profiles (Figure 4d) further illustrate the impact of sink variations along the phloem. For sink profiles with the highest demand at the top (‘LGSt’ and ‘LSt’), the transverse velocity is negative, indicating water movement from the phloem to the xylem. Conversely, for profiles with the highest demand at the base (‘LGSb’ and ‘LSb’), the transverse velocity is positive, signifying water movement from the xylem into the phloem. This behavior is governed by hydraulic equilibrium, where water moves from regions of higher water potential to regions of lower water potential to maintain balance across the vascular system. In the ‘LGSt’ and ‘LSt’ cases, sucrose removal near the top reduces osmotic potential at the source, driving water toward the xylem at the beginning of the phloem and into the phloem at the sink. On the other hand, for the ‘LGSb’ and ‘LSb’ cases, sucrose removal near the base sustains osmotic potential along most of the pathway, causing water to flow into the phloem at the source and toward the xylem at the sink. From an ecological perspective, the ‘LGSb’ and ‘LSb’ profiles represent more stable configurations. These profiles adhere more closely to the passive M’́unch mechanism, where sucrose transport is primarily driven by osmotically generated pressure gradients. In the current model, the imposed flux condition allows for both advection and diffusion of sucrose. However, under a fully passive transport regime (dominated by diffusion), maintaining an optimal osmotic potential gradient is essential for effective transport. This gradient naturally facilitates water movement into the phloem near the source and into the xylem near the sink, as observed in the ‘LGSb’ and ‘LSb’ profiles. This mechanism ensures efficient transport and ecological stability. Finally, the profiles of *v, u, p* (not shown), and *c* (not shown) are unaffected by the time of day (e.g., *t* = 24h or *t* = 16h). Only their magnitudes are influenced by external forcings such as fluxes and xylem water potential, as expected.

## 4. Conclusion

Mathematical modeling of phloem transport has been instrumental in advancing our understanding of the physical and physiological factors that govern sucrose movement in plants. However, simplifying assumptions, such as uniform sink profiles or constant viscosity, often limit the ecological and physiological relevance of these models. This study addresses these gaps by exploring the impact of stem sink variations on phloem dynamics using a detailed numerical model that incorporates transient dynamics and external forcings, such as xylem water potential and imposed fluxes. The results reveal that the spatial distribution of sucrose sinks along the phloem significantly influences flow speed, pressure gradients, and water exchange between the xylem and phloem. Notably, sink profiles with high sucrose demand near the base promote greater osmotic potential along the transport pathway, resulting in more stable and efficient transport dynamics. These profiles align with the passive M’́unch mechanism, wherein osmotic pressure gradients drive sucrose transport with minimal energetic cost. Conversely, sink profiles with high demand near the top reduce osmotic potential near the source, leading to lower pressure gradients and flow speeds. These profiles make the system more vulnerable to environmental stressors, such as reduced xylem water potential during drought or diurnal variations, which can compromise carbon allocation and transport efficiency.

A key finding of this study is that sink variations influence the hydraulic equilibrium between the xylem and phloem, as evidenced by transverse velocity profiles. For sink profiles with high demand at the top, water flows out of the phloem and into the xylem near the source, whereas the opposite occurs for profiles with high demand at the base. This behavior underscores the importance of osmotic potential gradients in maintaining a balanced hydraulic system and highlights how sink placement can enhance or hinder transport efficiency. From an ecological perspective, sink profiles with higher sink at the bottom represent a more resilient configuration, as they sustain osmotic potential throughout the pathway, ensuring robust transport even under fluctuating environmental conditions. The implications of these findings extend beyond individual plants to broader ecological and physiological contexts. Trees with sink profiles that promote gradual sucrose removal—expected to scale with the size of the stem—are better equipped to maintain efficient carbon allocation under environmental stress, such as drought or competition for light. Maintaining high osmotic potential near the source provides a critical buffer against adverse conditions, allowing trees to sustain effective sugar transport even when external stresses, such as low xylem water potential, challenge the hydraulic system. This mechanism may explain why certain tree species with similar physiological traits are able to thrive in environments characterized by high variability in water availability, as they can adapt their transport dynamics to meet changing demands. Additionally, the results offer new insights into the passive nature of phloem loading in large trees, particularly gymnosperms and some angiosperms. Previous studies have shown that these trees operate near the limits of efficiency under the M’́unch mechanism [8], and this study further emphasizes the importance of sink placement in optimizing transport dynamics. For instance, allowing small sink variations at the top can improve flow dynamics when sucrose transport relies on advection. However, this improvement comes at the cost of increased vulnerability to external forces, highlighting the trade-off between maximizing transport speed and maintaining system stability.

This work highlights the importance of incorporating spatial variations in sink demand into phloem transport models to capture the nuanced interplay between sucrose dynamics and plant physiology. Although the present study focuses on constant viscosity, accounting for concentration-dependent viscosity could uncover deeper interactions between sink profiles and viscous stress, offering a more complete picture of transport efficiency. The transient modeling approach used here opens the door to exploring how trees adjust sink distributions in response to environmental cues such as seasonal shifts or extreme events. Such adjustments likely play a vital role in enabling trees to maintain effective sugar transport and physiological stability under diverse and fluctuating conditions. These findings emphasize that sink variations are not just a passive aspect of phloem transport but an active determinant of transport efficiency, resilience, and carbon allocation. Understanding how trees dynamically regulate these processes provides critical insights into their adaptive strategies for surviving environmental stressors. Incorporating these dynamics into future studies and ecological models can deepen our understanding of long-distance transport in plants while improving predictions of their responses to challenges such as drought and climate change.

## Appendix A

### Numerical approximation using finite volume

The numerical method used in this study is based on the finite volume approximation, ensuring the conservation of mass. The domain is discretized into *n* grid cells, each of size Δ*z*, with conservation laws applied to each cell. The time evolution of mass (either sucrose or fluid) is determined by the difference between the inlet and outlet fluxes across the grid. To solve for the key variables, dynamic pressure *p*, sucrose concentration *c*, and dynamic viscosity *µ* are computed at the center of each grid cell, while the longitudinal velocity *u* is evaluated at the faces of the grid. Additionally, the transverse velocity *v* is explicitly solved, despite being expressible in terms of *u, p*, and *µ* due to its importance as a physical quantity in the problem. In this one-dimensional setup, *v* is solved at the center of the grid, reflecting its proportionality with *u*_*z*_. In a two-dimensional problem, however, *v* would be solved at the grid faces.

For the time evolution, Newton’s method for solving nonlinear PDEs is employed, utilizing an implicit time discretization scheme. This method efficiently handles non-linearities and provides fast convergence. The governing equations are presented in their nondimensional form, simplifying the analysis by focusing on key nondimen-sional numbers. This approach improves numerical behavior by scaling variables appropriately and reducing potential numerical errors.

#### Appendix A.1. Dynamic viscosity

The relationship between dynamic viscosity *µ* and sucrose concentration *c* is highly nonlinear [66], which makes the Jacobian matrices (discussed later) unsolvable analytically. Therefore, a simplification is introduced to achieve mathematical tractability. This is done using multiple linear regression to approximate dynamic viscosity in terms of sucrose concentration via a polynomial expansion:

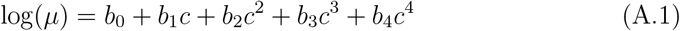

where the viscosity-concentration relationship from [66] is used to determine the coefficients *b*_0_, *b*_1_, *b*_2_, *b*_3_, *b*_4_. The equation is then nondimensionalized as follows:

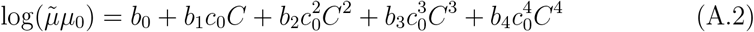

leading to

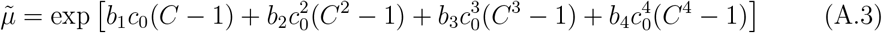

where *µ*_0_ is calculated from equation A.1 when *c* = *c*_0_. Finally, by introducing the relation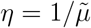, one can obtain the nondimensional dynamic viscosity equation:

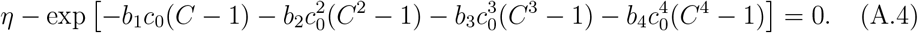

This new relation simplifies the derivation of the components of the Jacobian matrix, making the derivatives with respect to *C* more straightforward and easier to implement compared to the relation derived using a neural network [66].

The system of equations to be numerically solved is as follows:

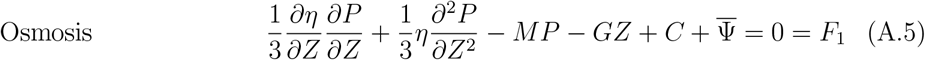

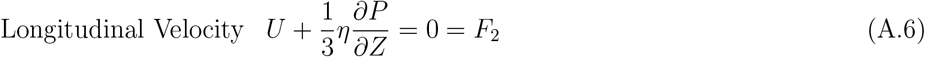

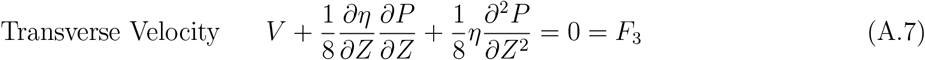

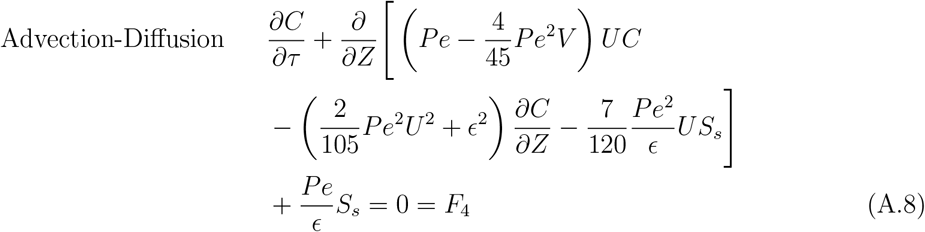

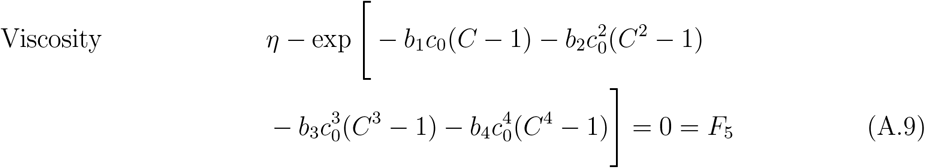

where *S*_*s*_ = *s*_*s*_(*c*_0_*u*_0_)^*−*1^ is a nondimensional number that quantifies the relative importance of the sink term and the advective term.

#### Appendix A.2. Newton’s method

Newton’s method is used to iteratively solve for ***S***^*j*+1^ given the nonlinear system of equations ***F*** (***S***^*j*+1^) = 0 where *j* denotes the time iteration. ***S*** is a vector of size (5*n −*1) *×* 1 that includes all the variables to be solved. These variables consist of *n* pressure points, *n −* 1 longitudinal velocity points (excluding the boundary points, which are specified separately), *n* transverse velocity points, *n* concentration points, and *n* viscosity points. The vector ***F***, also of size (5*n −* 1) *×* 1, contains the equations to be solved. This includes *n* equations of type *F*_1_ for pressure (osmosis), *n −* 1 equations of *F*_2_ for longitudinal velocity, *n* equations of type *F*_3_ for transverse velocity, *n* equations of type *F*_4_ for sucrose concentration, and *n* equations of type *F*_5_ for dynamic viscosity.

To solve the nonlinear system ***F*** (***S***^*j*+1^), Newton’s method linearizes the system around the current guess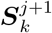(*k* denoting the iteration number before convergence) using a first-order Taylor expansion:

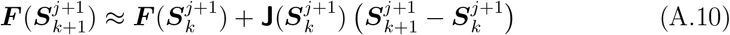

where **J** is the Jacobian matrix of size (5*n−*1)*×*(5*n−*1), which can be computed either analytically or numerically. In this case, ***F*** (***S***_*k*+1_) = 0 by definition, and at each iteration, one solves the system **J**_*k*_Δ***S*** = *−****F*** (***S***_*k*_) to compute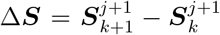which is then used to update the solution vector 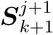. This process is repeated until convergence, meaning Δ***S*** is sufficiently small.

Due to the nature of the imposed boundary condition on *C*, which is a flux of sugar at *Z* = 0 and *Z* = 1 (Neumann boundary condition), the analytical Jacobian **J** becomes ill-conditioned. This implies that the system of equations may exhibit nearly singular behavior, leading to infinitely many solutions that depend heavily on the initial conditions. To ensure the stability of the numerical scheme, Tikhonov regularization is applied. Tikhonov regularization adds a regularization term to the Jacobian matrix to improve its conditioning and make it more suitable for inversion. The modified system of equations to be solved at each Newton iteration is (**J**_*k*_ + *λ***I**) Δ***S*** = *−****F*** (***S***_*k*_), where *λ* is a small positive regularization parameter, and **I** is the identity matrix of size (5*n −* 1) *×* (5*n −* 1). The regularization term *λ***I** helps stabilize the inversion of **J**_*k*_ by ensuring that the matrix is better conditioned. In this case, *λ* = 10^*−*6^ was selected after careful testing. The Euclidean norm of the residual error vector ***F*** (***S***_*k*_) was calculated to verify that this choice of *λ* sufficiently stabilized the system while maintaining solution accuracy.

#### Appendix A.3. Initial and Boundary Conditions

To solve the problem, boundary conditions must be specified for the pressure *p* at both ends of the phloem. Different formulations with varying physical interpretations can be chosen. For example, one could assume a closed tube at both ends where sucrose only enters the phloem through molecular diffusion (i.e., *u* = 0). However, due to the nature of the problem, where a sucrose flux is imposed, imposing a zero longitudinal velocity leads to a stiffer system and higher instability. For this reason, a zero inflow of water was chosen at both ends (*v* = 0). Physically, this condition implies that the xylem and phloem are at water potential equilibrium at the leaf and root levels. This assumption allows the system to maintain a balance in water flow and minimizes large gradients in pressure that could destabilize the solution. Additionally, the sucrose flux at the leaf level, *s*_*l*_, was chosen to balance the sink at the stem and the root (see Table 1). In this setup, during each forcing time step (i.e., during the period where externally imposed variables such as stem sink and xylem water potential from BiomeE remain constant), the sucrose mass within the phloem is conserved. In this case, phloem hydrodynamics are more sink-limited.

Since the initial conditions are unknown and the model is solving phloem hydrodynamics with an imposed forcing from BiomeE, two possible initial conditions were considered. The first approach assumes that the initial osmotic profile is proportional to the hydrostatic and xylem potential, given by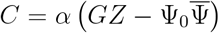, where *α* is a constant. This approach allows the study of the difference between steady-state and transient phloem conditions, especially during times when there is a significant offset between the forcing variables and phloem dynamics. For the results in Section 3, *α* = 2 was chosen. The second approach assumes the phloem is already in a steady-state condition within the plant, meaning the same initial guess is used, but Newton’s iterative method is applied with *∂C/∂τ* = 0. This approach is useful for cases where the phloem system is assumed to have reached equilibrium prior to the application of external forcing, which is physically accurate since the entire plant system is assumed to be evolving in a balanced manner.

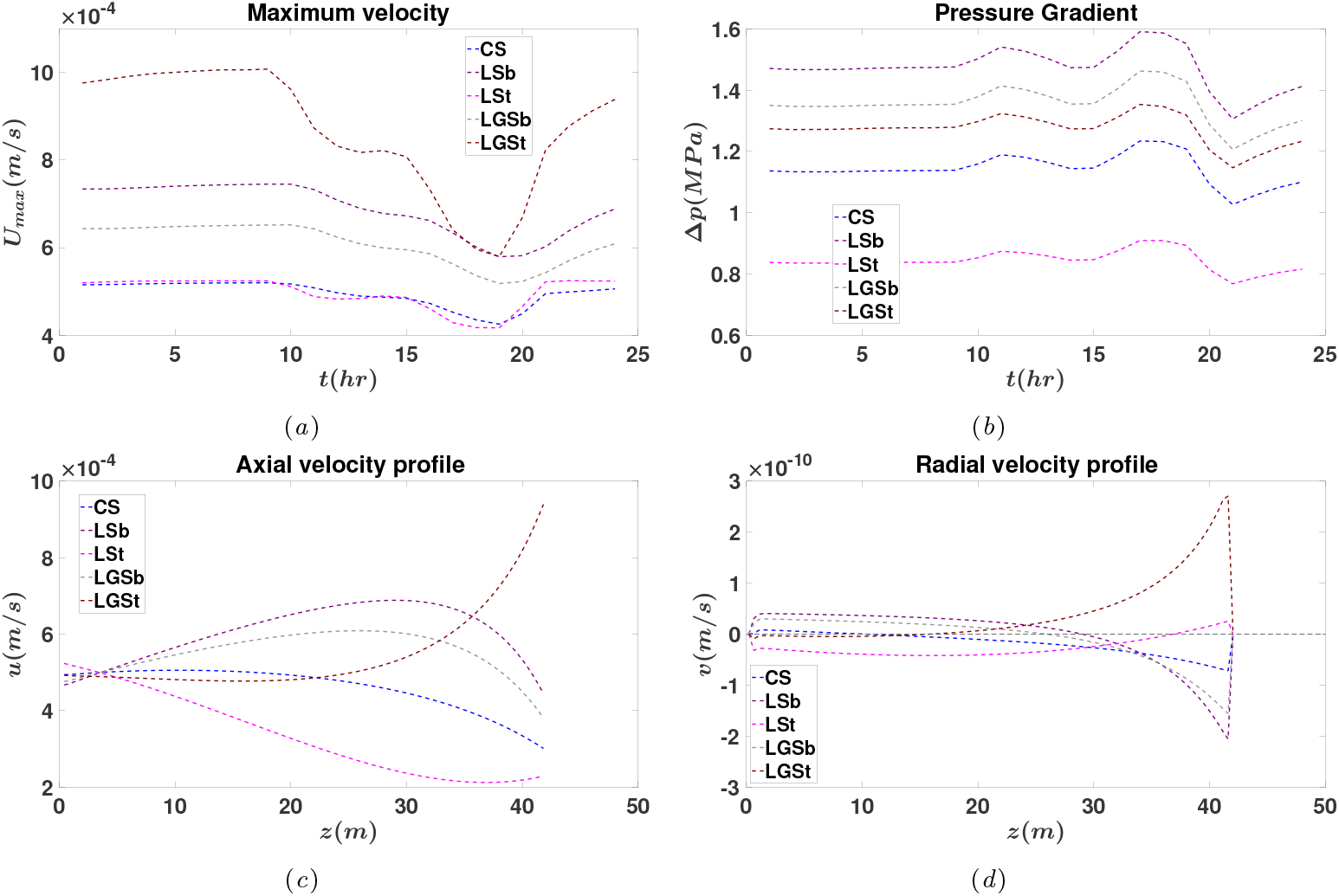

